# The range-resident logistic model: a new framework to formalize the population-dynamics consequences of range residency

**DOI:** 10.1101/2025.02.09.637279

**Authors:** Rafael Menezes, Justin M. Calabrese, William F. Fagan, Paulo Inácio Prado, Ricardo Martinez-Garcia

## Abstract

Individual movement is critical in shaping population dynamics. However, existing frameworks linking these two processes often rely on unrealistic assumptions or numerical simulations. To address this gap, we introduce the range-resident logistic model, an easy-to-simulate and mathematically tractable extension of the spatial logistic model that incorporates empirically supported range-resident movement. Our framework unifies non-spatial and (sessile) spatial formulations of the logistic model as limiting cases. Between these regimes, the long-term population size depends nonlinearly on home-range size and spatial distribution. Neglecting range residency can hence lead to under- or overestimating population carrying capacity. To better understand these results, we also introduce a novel crowding index that depends on movement parameters and can be estimated from tracking data. This index captures the influence of spatial structure on population size, and serves as a robust predictor of abundance. The range-resident logistic model is thus a unifying framework bridging movement and population ecology.

## 1 Introduction

Recent advances in tracking technology, data-sharing platforms, and statistical analyses have provided unprecedented insight into how organisms move, including a more precise quantification of their home ranges (Kays *et al*. 2015; Kranstauber *et al*. 2011; Langley *et al*. 2024; McClintock *et al*. 2020; Silva *et al*. 2022). Home ranges, which encapsulate the areas that animals regularly use for securing resources and caring for offspring, are ubiquitous across animal species (Burt 1943). Home-range size and overlap determine how often organisms encounter one another, which makes them key drivers of intra and interspecific interactions (Berger & Gese 2007; Fagan *et al*. 2024; Martinez-Garcia *et al*. 2020; Powell 1979). However, although we can accurately estimate home-range sizes as well as encounters and their consequences for range-resident animals, we still do not fully understand their quantitative impact on population and community dynamics (Ayllón *et al*. 2012; Doherty 1983; Doncaster 1990; Fagan *et al*. 2024; Fieberg & Kochanny 2005; Hartmann *et al*. 2017; Knapton & Krebs 1974; López-Sepulcre & Kokko 2005; Winner *et al*. 2018). Establishing a theoretical link between range-resident movement and population dynamics is critical to answering many applied questions in population and community ecology (Costa-Pereira *et al*. 2022; Morales *et al*. 2010).

Theoretical ecology has mainly developed from non-spatial models that only track abundances and assume homogeneous distributions of populations where organisms interact via the law of mass action (Hutchinson & Waser 2007; Lotka 1920; Verhulst 1838; Volterra 1926). This approach oversimplifies movement and overlooks the impact of within-range movement on population dynamics. Alternative approaches, such as metapopulation models or reaction-diffusion equations, incorporate different movement and dispersal modes (Cantrell & Cosner 2004; Hanski 1998; Levins 1969). However, they still assume well-mixed conditions at the scale of interactions. Other models incorporate movement implicitly, combining the simplicity of non-spatial models with phenomenological descriptions of territorial behavior to link carrying capacities with territory sizes and environment quality (Ayllón *et al*. 2012; López-Sepulcre & Kokko 2005). Despite their merits, these models are insufficient to describe the more general case of non-territorial populations with varying levels of home range overlap. Finally, individual-based models (IBM) can describe movement at almost any level of detail, including range residency, and might be easier to parameterize for specific populations (Grimm & Railsback 2013). Computational simulations of IBMs with range-resident movement have helped us understand the causes of territoriality (Giuggioli *et al*. 2011), and how environmental and intraspecific trait variability promote the stability of animal communities (Buchmann *et al*. 2011, 2012; Milles *et al*. 2020).

Therefore, IBMs have revealed some of the mechanisms underlying home-range spatial dynamics and are more straightforwardly transferred to conservation and management. Yet, with current analytical tools, IBMs are mathematically tractable only for Brownian movement—which does not describe range residency and converges to well-mixed conditions if organisms explore the entire available environment between demographic processes—or sessile organisms (Bolker & Pacala 1997; Dieckmann *et al*. 2000; Plank & Law 2015). This lack of mathematical tractability and their system-specific nature make most IBMs an inappropriate basis from which to extract general conclusions on how movement parameters affect population dynamics. A promising direction lies in identifying alternative stochastic processes that can simultaneously maintain analytical tractability while improving the realism of simulated animal movement. The Ornstein-Uhlenbeck (OU) stochastic process is an optimal compromise, as it accurately captures the two main features of range residency: home ranges covering only a fraction of the total population range and non-uniform utilization of each home-range area. Additionally, recent multi-species comparative studies of home ranges based on GPS tracking data identified OU process and its variant incorporating autocorrelated velocities—the Ornstein-Uhlenbeck with Foraging (OUF) process—as the AIC-best models with which to perform home-range estimation in 368 out of 369 (Noonan *et al*. 2019), and 1235 out of 1239 (Fagan *et al*. 2025) datasets. Finally, the OU stochastic process has a long tradition in ecology (Dunn & Gipson 1977) and can be readily extended to account for more complex animal behaviors (Blackwell 1997; Blackwell *et al*. 2016; Smouse *et al*. 2010), including switching among multiple home-range centers (Breed *et al*. 2017) or resource selection (Eisaguirre *et al*. 2021).

Ideally, a general theory to investigate the population-level consequences of non-territorial range residency should recover a description of sessile organisms in the limit of vanishingly small home ranges and converge to nonspatial frameworks as home ranges increase in size and organisms explore them faster. Additionally, starting from an individual-level description of how organisms move and interact in space, this general theory should provide a dynamical equation for population abundance whose parameters contain information about home-range size and overlap. Encounter rates are the bridge between movement and population dynamics, and thus provide a good starting point to upscale the consequences of movement to population dynamics. The first steps toward such a framework have focused, therefore, on deriving temporal and spatial statistics for the encounters between pairs of organisms with partially overlapping home ranges (Martinez-Garcia *et al*. 2020; Noonan *et al*. 2021). Moreover, these first results already show that range-resident encounter rates can deviate significantly from law-of-mass-action expectations (Martinez-Garcia *et al*. 2020). Yet, a complete upscaling embedding non-territorial range-resident interactions into a birth/death population dynamics framework is still to be performed.

Here, we develop the range-resident logistic model as a unified spatial modeling framework linking individual movement behavior to its population-level consequences. In particular, the range-resident logistic model allows us to quantify how interactions between home-range size, spatial distribution, and intraspecific competition affect population growth and carrying capacity. The proposed model utilizes the OU stochastic process to represent movement, capturing the key features of range-resident movement while being supported by tracking data and maintaining mathematical tractability. We also demonstrate how the range-resident logistic model includes both the classical logistic and spatial logistic models in the limits of very large and vanishingly small home-range size, respectively. Importantly, we show that assuming well-mixed conditions may either under- or overestimate population abundances. This new range-resident logistic framework thus unifies existing population dynamics models, providing important insights into how individual movement influences patterns observed at higher levels of organization.

## 2 Materials and Methods

### 2.1 The range-resident logistic model

We study a single-species population inhabiting a homogeneous habitat patch of area *A* and model its dynamics using an individual-based approach. The model incorporates different movement behaviors throughout the organism’s lifetime and includes the minimum set of stochastic demographic processes required for density-dependent population growth: density-independent reproduction followed by dispersal and density-dependent death driven by intraspecific competition (Fig. 1). Apart from introducing range-resident movement, the model closely follows well-established logistic IBMs (Bolker & Pacala 1997; Law *et al*. 2003).

**Figure 1:**
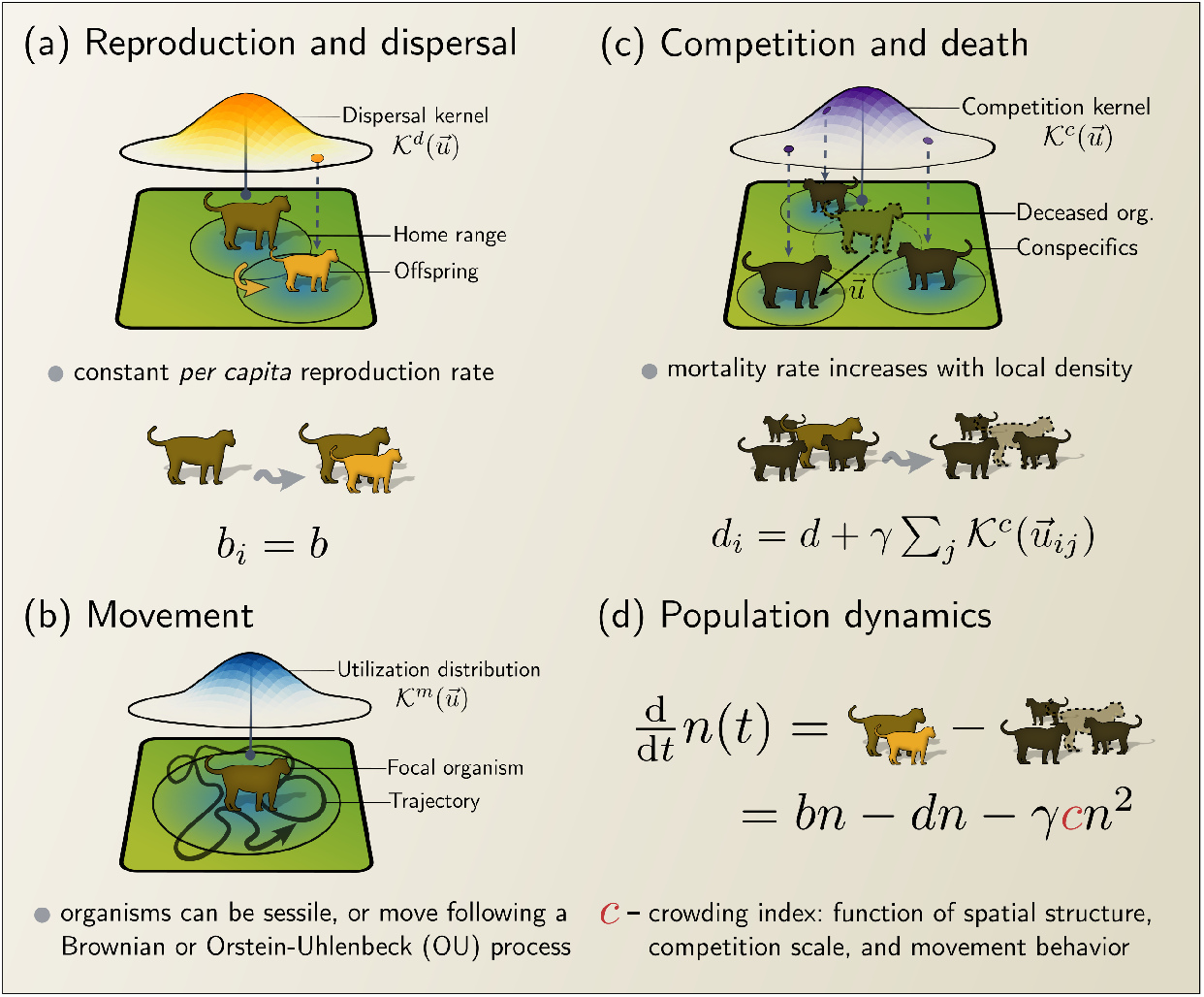
Graphical summary of the range-resident logistic model. (a) Organisms reproduce at a constant rate *b* with offspring establishing new home ranges at a position sampled from a juvenile dispersal kernel, 𝒦^*d*^(***u***), centered at the parent’s home range center. (b) Organisms move following stochastic movement models. After each simulation step, the position of the organisms were updated by integrating the corresponding stochastic differential equation. (c) Organisms experience a baseline mortality rate of *d* that grows linearly with the local density of organisms. This local density around each organism is calculated using a competition kernel 𝒦^*c*^(***u***). (d) Population abundance changes over time due to the balance between birth and death. Spatial factors such as movement and dispersal change the aggregation pattern, measured by the crowding index *c*.

#### Movement

We assume that changes in animal location occur on a much faster timescale (hours) than demographic processes (months to years), and animal trajectories are given by a continuous-in-time realization of a stochastic process (Fig. 1B). Under this assumption, movement is independent of any demographic process, and we can update the position of every organism in the population after each demographic event by integrating the stochastic differential equation that describes the movement process for each organism. Although our focus is on how range residency influences population dynamics, we do not limit our analyses to range-resident movement. We consider:

‐ *Range-residency*. We use the Ornstein-Uhlenbeck (OU) process, a data-supported (Noonan *et al*. 2019) and well-established stochastic model for range-resident movement. The OU process is a drift-diffusion process in which random, diffusive movement is complemented by the deterministic tendency of range-resident animals to return to the vicinity of a preferred location (home-range center). This combination of processes leads to home ranges smaller than the population range and non-uniform space use within each home range (Blackwell 1997). Assuming that movement is two-dimensional, isotropic, and uncorrelated in each of its components, trajectories of the OU model are generated by the following Langevin equation:

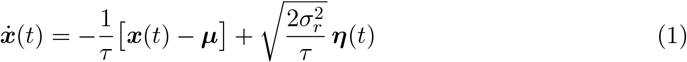

where ***x*** and ***µ*** are two-dimensional vectors containing the organism and its home-range center locations, respectively; *τ* is the average home-range crossing time; 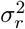 defines the characteristic spatial scale of the home-range size by setting the long-term variance of the animal’s space utilization function, and ***η***(*t*) is a two-dimensional white noise vector with zero-mean and unit-variance components. At sufficiently long time scales, the individual space utilization functions that result from the OU movement in Eq. (1) are isotropic Normal distributions with mean ***µ*** and variance 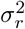. Equivalently to the trajectory-based description via a stochastic differential equation, the Ornstein–Uhlenbeck process can be formulated as an advection–diffusion (Fokker–Planck) equation for the probability distribution of the animal’s location. In this representation, diffusion is 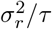 and the advection velocity (***x***−***µ***)*/τ* (Smouse *et al*. 2010). This description is closer to a mechanistic home-range analysis framework (Moorcroft & Lewis 2013), but it is less common in individual-based models of population dynamics (Plank *et al*. 2025).
‐ *Uniform space usage*. To turn off the effect of range residency on the observed patterns of population dynamics, we also consider the simpler case in which organisms do not exhibit range residency and hence explore the whole population range. We can recover this movement behavior from Eq. (1) in the limit in which *τ* → ∞ and 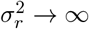 with the diffusion coefficient 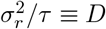 finite (Martinez-Garcia *et al*. 2020).
‐ *Sessile*. Finally, we also consider the limit in which organisms do not move during their lifetime and thus remain where they land upon dispersal. In terms of parameter values, this limit corresponds to taking *τ* → ∞ and 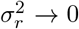 with the diffusion coefficient *D* = 0. In this limit, our model is equivalent to the spatial logistic model introduced by Bolker & Pacala (1997), and Law *et al*. (2003).

#### Demographic processes

‐ *Density-independent reproduction and dispersal*. Organisms in our model have a constant intrinsic birth rate *b* (Fig. 1A) and they disperse and establish their own home-range centers immediately upon birth. We sample the dispersal displacement between the offspring’s home-range center and their parent’s, ***u***, from a dispersal kernel, 𝒦^*d*^(***u***). We use a bivariate Normal distribution centered on the parent’s home-range center with standard deviation *σ*_*d*_ that sets the characteristic dispersal distance (Fig. 1A). Hence, the distance between the parent’s and offspring’s home-range centers follows a Rayleigh distribution with non-zero mode. For range-resident organisms, we additionally sample the initial position of the offspring from its asymptotic space utilization distribution, assuming that organisms relax to their steady-state movement behavior immediately after dispersal.
‐ *Density-dependent death*. We consider that organisms die at a rate that depends on how crowded their local environment is. For a focal organism *i*, its death rate is a function of its position ***x***_*i*_, and is given by

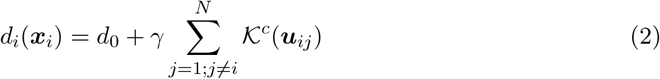

where *γ* is the competition strength, *N* is the total number of organisms in the population, *d*_0_ is the baseline death rate, and 𝒦^*c*^(***u***_*ij*_) is a competition kernel defining how competition intensity decays with the displacement between the focal organism *i* and its competing neighbor *j* (Fig. 1C). For the competition kernel, we choose a bivariate normal distribution (and thus normalized to one) with standard deviation *σ*_*q*_, centered at the position of the focal organism *i* (Law *et al*. 2003).

### 2.2 Analytical tools and approximations

#### 2.2.1 Spatial moment dynamics equations

Spatial moment dynamics (SMD) is a mathematical method to approximate the dynamics of spatially explicit IBMs by a hierarchy of coupled ordinary differential equations describing the dynamics of the moments of the spatial distribution of organisms (Iwasa 2000; Markham *et al*. 2013; Murrell *et al*. 2004; Plank & Law 2015; Simpson *et al*. 2014). This hierarchy of equations is often truncated at the second order, which accounts for the coupled dynamics of the mean density of organisms (first moment) and the mean density of organism pairs separated by a displacement vector ***u*** (spatial correlations; second moment). The second-order SMD equations are hard to treat when home-range centers are organism-specific. Alternatively, we work at first order and quantify spatial structure in terms of a crowding index that depends on movement and dispersal parameters and also retains information on the local population density each organism experiences (Wiegand *et al*. 2021).

To obtain the first-order SMD equation, we need first to derive the continuous equivalent of the discrete rates introduced in the IBM dynamics. The per-capita birth rate is assumed constant and equal to *b*, whereas the per-capita death rate increases with local crowding accounting for intraspecific competition. To obtain the average death rate in the population, we need to compute the probability that a focal organism has a neighbor at a displacement ***u***. These spatial correlations are provided by the population pair correlation function, *g*_2_(***u***, *t*), which is a dimensionless second-order spatial metric measuring the ratio between the density of organism pairs separated by a displacement ***u*** and the average density of pairs, *n*^2^ (Baddeley *et al*. 2016; Law *et al*. 2009; Wiegand *et al*. 2025)—where *n* is the density of organisms *n* = *N/A*. Therefore, *g*_2_(***u***, *t*) is constant and equal to one in homogeneous populations, while values greater or lower than unity indicate aggregation or overdispersal at a specific displacement ***u***. In terms of this metric, an arbitrary organism has an average number of neighbors at displacement ***u*** equal to *ng*_2_(***u***) (see e.g., Plank & Law 2015, for detailed derivations), and the continuous per-capita death rate is

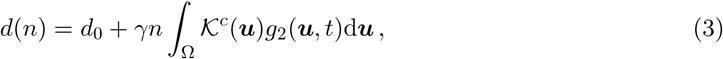

where the integral is taken over the domain Ω of all possible displacements between organisms.

Under our assumption of no net immigration or emigration, movement does not directly contribute to population size, and hence, the dynamics of the first spatial moment only depends on the balance between the birth and the death rate. Using the expressions for the per-capita birth and the death rate in Eq. (3) we can write the dynamical equation for the density of organisms as

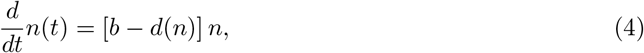

which recovers the classic logistic equation when individuals are uniformly distributed and, consequently, *g*_2_(***u***, *t*) = 1.

#### 2.2.2 Crowding Index

Two key functions, the pair correlation function *g*_2_(***u***, *t*) and the competition kernel 𝒦^*c*^(***u***), are central to understanding how population abundance depends on spatial scale. The pair correlation function serves as a geometrical aggregation index characterizing the typical distance between organisms. In contrast, the competition kernel specifies how the strength of the competitive interaction changes with the spatial scales. The integral involving these two functions in Eq. (3) indicates that the death rate depends on the local distribution of organisms within the spatial scales at which they interact, but is insensitive to how organisms are distributed at other scales. To convey this information, we introduce the crowding index *c*(*t*)

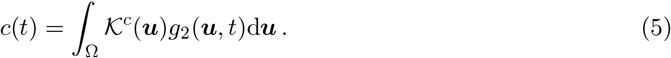

which is a weighted second-order spatial statistic using the competition kernel 𝒦^*c*^ as a weighting function.

The definition of the pair correlation function and the normalization of the competition kernel set a reference value for the crowding index, *c*(*t*) = 1, in homogeneous populations. Index values greater than one indicate some degree of overcrowding at the scales at which individuals compete, and more intense competition relative to homogenously distributed organisms. Conversely, *c*(*t*) *<* 1 indicates undercrowded populations, and organisms interacting less often than in the homogeneous setting. Note also that the crowding index incorporates information across the entire interaction range, and thus a “surplus” of neighbors at one scale might offset a “deficit” at others.

This interpretation of the crowding index, which emerges directly from the demographic rates, is equivalent to other definitions using the weighted covariance (Bolker & Pacala 1997, 1999). The only difference is that we use the pair correlation function *g*_2_(***u***, *t*) as the second-order spatial statistic instead of using the spatial covariance ⟨*n*(***x***) − *n, n*(***y***) − *n*⟩, where *n*(***x***) is the local density of organisms at the position ***x***. Using the pair correlation function instead of the spatial covariance avoids the introduction of self-competition terms in the resulting population dynamics. The crowding index in terms of the weighted covariance, *c*_cov_, is related to the crowding index defined in Eq. (5) by *c*_cov_ = *n*^2^(*c* − 1).

### 2.3 Numerical simulations

We simulated the IBM of Section 2.1 using Gillespie’s algorithm (Gillespie 1977), which we implemented in Python3 using the Numpy, Scipy, and Numba libraries (Harris *et al*. 2020; Lam *et al*. 2015; Virtanen *et al*. 2020). All source code, original data, and scripts required to reproduce the individual-based simulations, analyses, and figures for the main text and supplementary material are archived in the Zenodo repository at https://doi.org/10.5281/zenodo.15312822.

We performed all simulations in a square domain of lateral length *L* = 1, using periodic boundary conditions and setting the reproduction, basal mortality, and interaction rates at *b* = 1.5, *d* = 0.1, and *γ* = 0.002, respectively. Other parameters were varied depending on the analyses, and their values are indicated in figure captions and legends. We truncated the competition kernel at 2.45 times the standard deviation, capturing 95% of its probability mass. For each specific parameterization, we ran 20 independent replicates, using a spatially uniform distribution of 700 organisms as the initial condition, which corresponds to the carrying capacity of the non-spatial model, 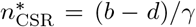. Each of these replicates ran for 10^6^ simulation events, after ensuring, by visual inspection, that the population had reached a quasistationary state (see Fig. C.1 for time series of population abundance in some representative scenarios). At the end of each simulation run, we recorded the total abundance and the spatial distribution of organisms, which we used to compute statistical metrics (medians, means, and percentile intervals). Finally, we normalized the population carrying capacity recorded in the simulations, *n*_*t*→∞_, dividing it by 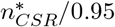 to account for the reduced effective interaction rate induced by the truncation of the competition kernel. This normalization allowed for a clear assessment of how the carrying capacity of the range-resident logistic model deviates from the homogeneous case.

## 3 Results

### 3.1 Numerical simulations of the range-resident logistic IBM

We first performed numerical simulations of the range-resident logistic IBM to understand how movement and spatial processes determine carrying capacity. Home-range size modulates the population carrying capacity, with larger home ranges leading to carrying capacities closer to that predicted by the non-spatial model (Fig. 2). For very large home ranges, organism movement is effectively Brownian (BM), with home ranges covering almost the entire simulation domain. In this limit, the population is homogeneously distributed in space, consistent with the non-spatial (or well-mixed) limit assumptions. For vanishingly small home ranges, organisms are effectively sessile (SS) and located at their home-range centers, with interaction rates depending only on the distance between home-range centers. Organisms whose home-range centers are close to each other interact much more strongly than pairs of organisms whose home-range centers are far away. Therefore, in this limit, the carrying capacity is strongly determined by the characteristic dispersal distance, *σ*_*d*_: short-range dispersal led to population collapse, while the total abundance for long-range dispersal was 75% larger than that of homogenous populations (Fig. 2). Interestingly, the carrying capacity exhibited a minimum when home range, competition, and dispersal spatial scales were similar to each other, highlighting a non-trivial relationship between carrying capacity and home-range size (orange line in Fig. 2; supplementary figure C.2). This non-monotonic relationship is robust against changes in the dispersal kernel, as confirmed by an additional analysis using Gamma-distributed dispersal distances (see supplementary figure C.3).

**Figure 2:**
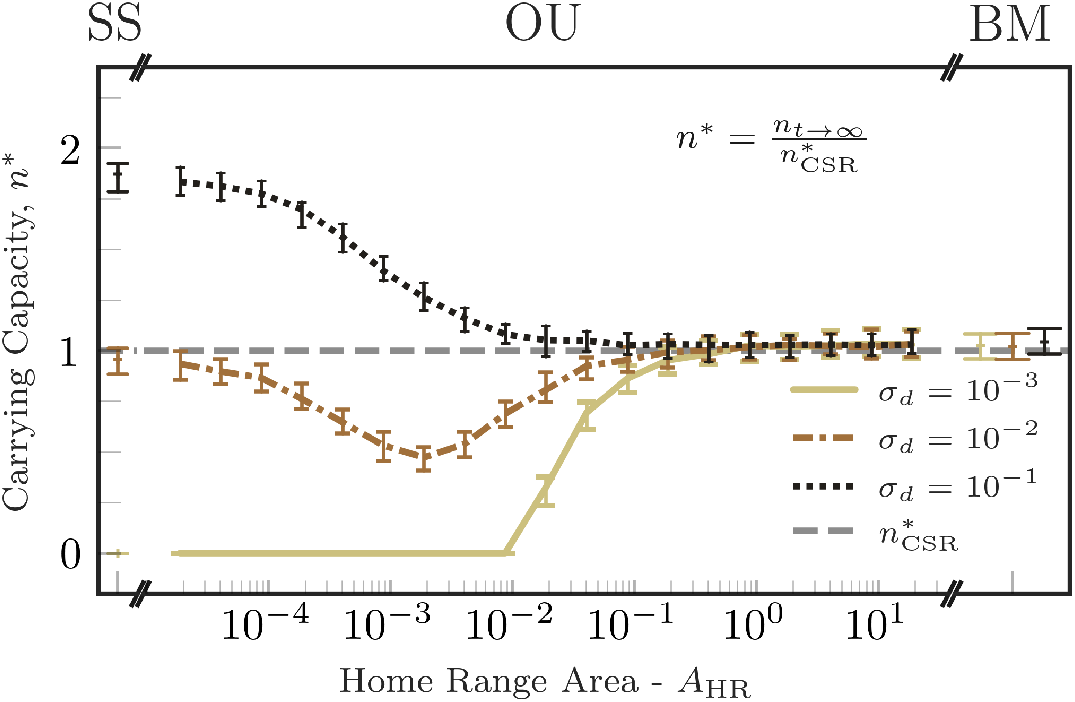
Home-range size is a key driver of the population carrying capacity. The carrying capacity, measured directly from IBM simulations and scaled by the carrying capacity of the non-spatial model 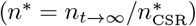 (gray-dashed line), versus home-range size. Different colors represent different dispersal ranges, as indicated in the legend. Curves and vertical bars correspond to the median and 90% percentile interval of the carrying capacity measured across realizations with the same parameter set. The labels on the top indicate the movement model used in the simulations: sessile (SS), range-resident movement (Orstein-Uhlenbeck; OU), and uniform space use (Brownian movement; BM). For OU movement, we changed the home-range area by increasing home-range crossing times while keeping diffusion fixed. When the home-range area and dispersal were small, populations go extinct.

Given the complex dependence of the carrying capacity on the various model parameters, we performed extensive simulations varying home-range size and crossing time, and competition and dispersal ranges over several orders of magnitude (Fig. 3). The competition range defines the scale at which organisms interact with each other, and thus has a strong impact on the carrying capacity. Long-range competition results in carrying capacities similar to that of the non-spatial model (rightmost column in Fig. 3). Short-range competition, however, leads to highly variable values of the carrying capacity (darker colors in the leftmost column of Fig. 3). Increasing the dispersal range generally leads to higher carrying capacities because offspring are more likely to establish new home ranges farther from their parents, which both avoids direct competition with them and increases the proportion of the available habitat the population colonizes (rows from bottom to top in Fig. 3). Finally, the carrying capacity is insensitive to changes in the home-range average crossing time, except at very large values of *τ* for which the carrying capacity increases quickly with *τ* (y axes in Fig. 3). In this limit, organisms move very slowly and only explore a small fraction of their asymptotic home ranges before dying (see supplementary figure C.4). In contrast, when the home-range average crossing time is very short compared to the temporal scale of demographic dynamics, individual positions relax to their stationary probability distribution function between demographic events.

**Figure 3:**
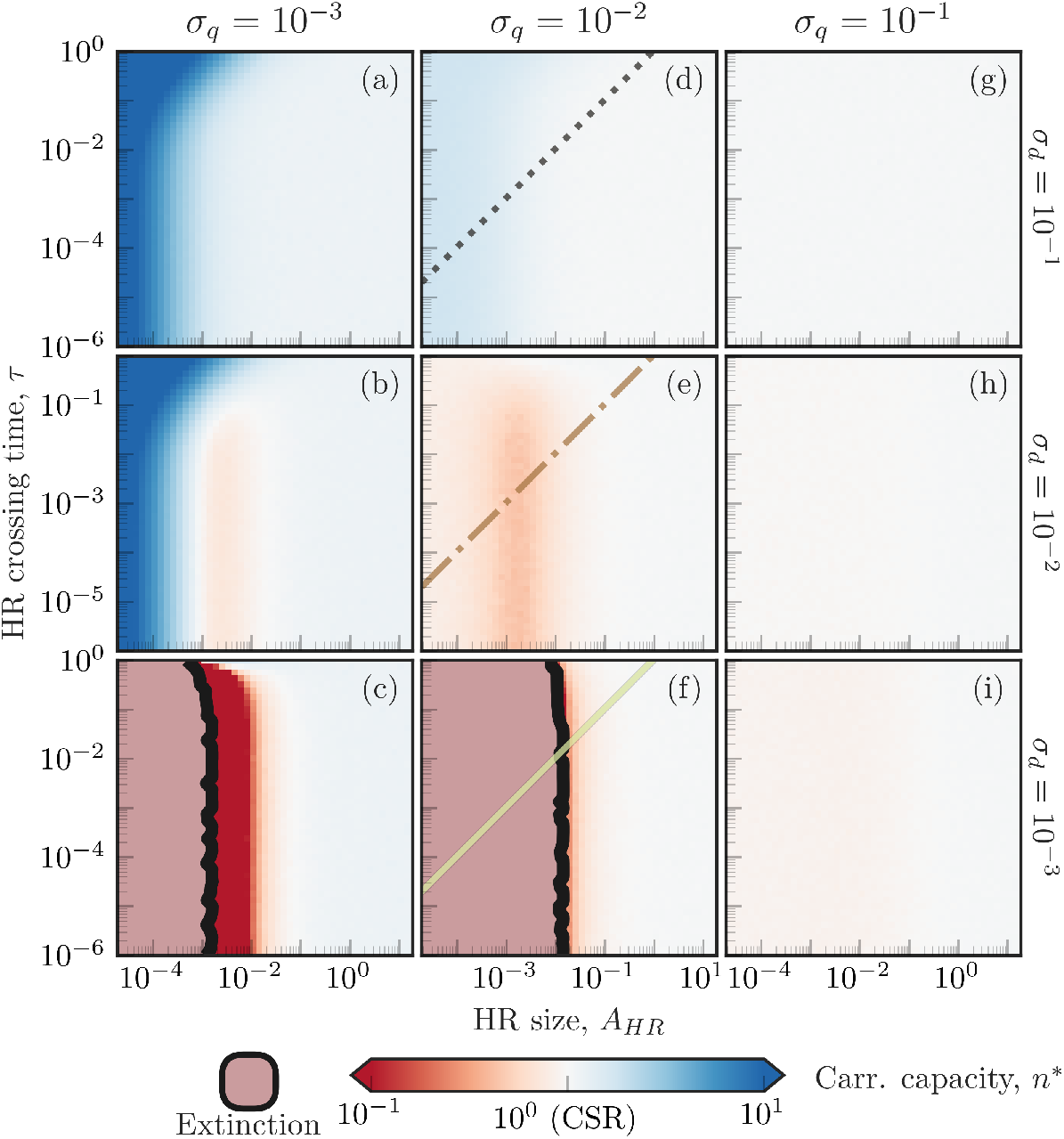
Competition and dispersal scales change the carrying capacity of the population across varying home-range (HR) sizes and crossing times. Across all panels, the carrying capacity of the population measured from simulations 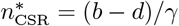 is plotted against the HR size (bottom axes), the HR crossing time (left axes), the competition scale (columns; top axis), and the juvenile dispersal scale (rows; right axis). Increasing HR size or competition range leads effectively to non-spatial dynamics, regardless of the value of other model parameters (lighter colors in the heatmaps). The HR average crossing time *τ* does not change the carrying capacity, except for large *τ* and small HRs where organisms are effectively sessile. Increasing juvenile dispersal allows for larger population abundances across all the parameter ranges considered. The colored lines in the central column indicate the parameter combinations used in Fig. 2, following the same color code used there.

### 3.2 Formulas relating crowding, home-range overlap, and population size

To understand the relationship between the population carrying capacity and individual movement behavior, we constructed a first-order SMD approximation of the IBM and analyzed its equilibrium population sizes. As we showed in the Methods, the dynamics of the total density of organisms is given by

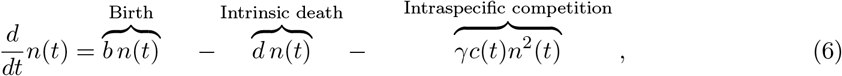

where *c*(*t*) is the crowding index at time *t* defined in Eq. (5). If the population is homogeneously distributed in space, *c*(*t*) = 1 and the non-spatial logistic model is recovered (Verhulst 1838). Imposing 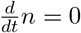 in Eq. (6) and solving for *n* returns two possible equilibria. The first one, 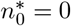, indicates population extinction. The second one is a non-trivial solution that defines the carrying capacity

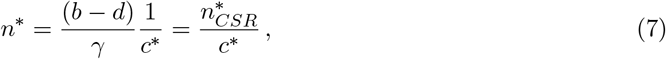

where *c*\* is the expected crowding index once the population reaches equilibrium. Equation (7) indicates that the ratio between the carrying capacities of the range-resident logistic model and the non-spatial model (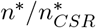, depicted in Figs. 2–3) is equal to the inverse of the crowding index at equilibrium. The competition, dispersal, and movement spatial scales will thus affect the carrying capacity by modulating the population spatial distribution and hence changing crowding intensity. Each of these scales, however, plays a different role in defining crowding intensity. The competition kernel 𝒦^*c*^ is a weighting factor accounting for how strongly a given spatial configuration of organisms impacts the survival of a focal organism. Dispersal and movement behavior define the population spatial structure and, consequently, change the crowding index by altering the pair correlation function of organisms’ positions *g*_2_(***u***, *t*).

To explicitly investigate how crowding depends on movement behavior, we redefine the crowding index in terms of the size and distribution of home ranges in space. Thus, we calculate the crowding index based on the pair correlation function of the point pattern of organisms’ home range centers 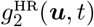. In this case, the weighting function will depend both on the competition kernel and on the pattern of space usage of each organism around its home range center, which we assume to be constant across the population in our simulations. Denoting the weighting function for the crowding of home ranges as 𝒦^HR^(***u***), it follows that

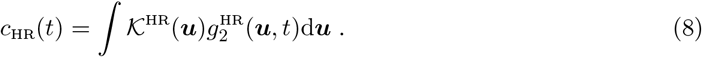

For the common case in which range-resident animals have mean home-range crossing times much shorter than the demographic time scales, i.e. *τ* ≪ *b*+*d*, Eq. (8) is equivalent to (5) (see supplementary material A) and our range-resident logistic model is equivalent to a version of the (sessile) spatial logistic model with modified competition kernels (Bolker & Pacala 1997; Law *et al*. 2003). Specifically, if interactions among organisms are local, i.e. 𝒦^*c*^(***u***) → *δ*(***u***), the range-resident logistic model is equivalent to a model of “sessile organisms” (the home range centers) interacting via a competition kernel that is defined by the stationary distribution of displacements between organisms (i.e., the distribution of displacements between organisms when organism locations are sampled from the space utilization distribution, see supplementary material A). This relationship thus provides a mathematical expression relating crowding and, consequently, the total population size to home-range overlap.

### 3.3 Crowding indexes as predictors of total population size

Finally, we tested the accuracy of the crowding index in predicting the carrying capacity of the population. From equation (7), the inverse of the crowding index is equal to the normalized carrying capacity, and thus provides a way to anticipate the population carrying capacity using only spatial information. We measured the crowding index using the competition scale and either information about organism locations or HRs’ centers and size for a large subset of simulations (Fig. 3 panels d-i; see supplementary material B). We compared the normalized carrying capacity measured in the simulations with the inverse of the measured crowding index in a predicted versus measured diagram (Fig. 4). We initially did not consider small interaction scales in these results, (*σ*_*q*_ = 10^−3^, Fig. 3 panels a-c) because local interactions are associated with very large effective interaction rates when organisms are close together and thus introduce deviations from our theory that considers mean interaction rates (Das *et al*. 2023; de Figueiredo *et al*. 2025). The predicted and measured carrying capacities showed a remarkable similarity (*R*^2^ = 0.983 for 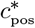 and *R*^2^ = 0.987 for 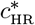). Including small interaction scales in the analysis decreased the accuracy of predictions, but the crowding index still explained most of the variance in the data (*R*^2^ = 0.893 for 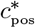 and *R*^2^ = 0.827 for 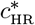, supplementary figure C.5). These results indicate that the crowding index, even when computed entirely using movement parameters that describe long-term animal movement, is a reliable predictor of the carrying capacity of the population.

**Figure 4:**
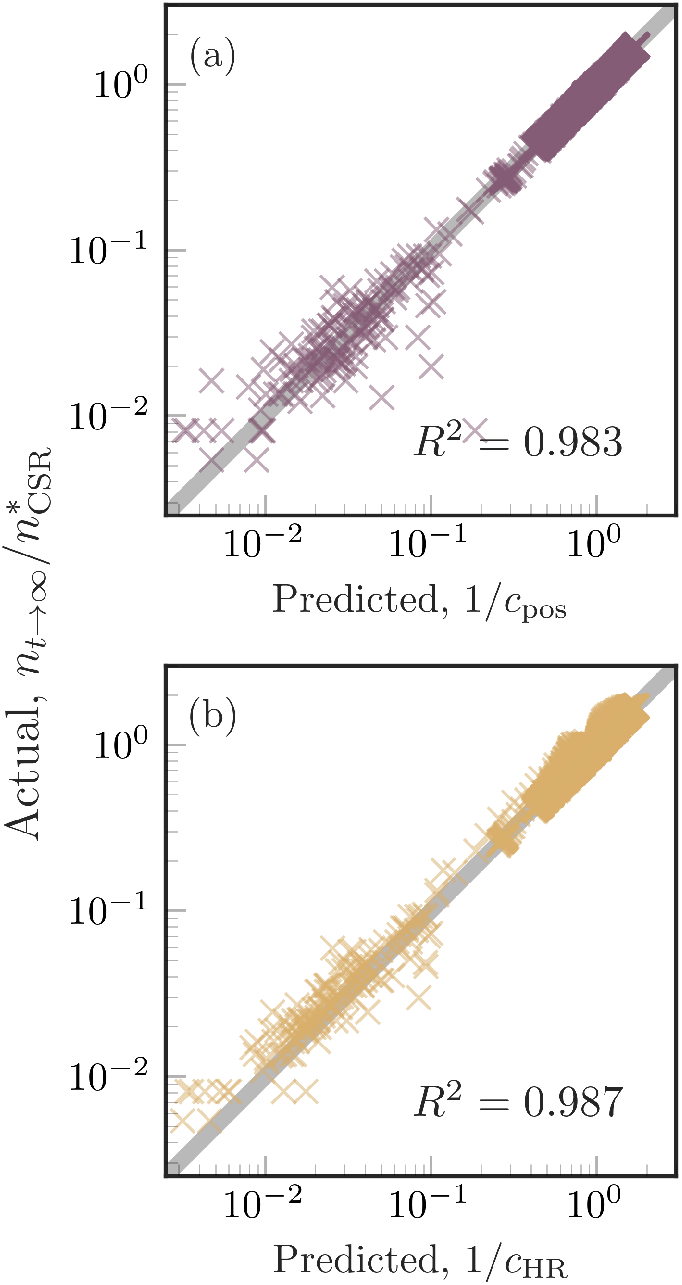
Crowding indexes accurately predict population carrying capacity. (a) The crowding index measured using organism locations in simulation snapshots. (b) The crowding index measured based on the expected mortality rate an organism experiences given its home-range size and distances between home-range centers. Gray lines in the background of each panel represent the complete agreement (1-1 line), and *R*^2^ values are provided as indicators of the goodness of the linear fit.

## 4 Discussion

We generalized existing spatial logistic models (Bolker & Pacala 1997; Law *et al*. 2003) to incorporate two key features of range-resident movement: organisms moving within home ranges that are smaller than the population range and exhibiting uneven patterns of space use. These two features shape a tradeoff in how organisms interact: bigger ranges provide more opportunities to interact with neighbors, but each individual competitor is encountered less often and with weaker effects (Fig. 5). This tradeoff is particularly relevant when home-range sizes are comparable to the spatial scales of dispersal and competition, possibly leading to non-monotonic relationships between home-range size and carrying capacity, consistent with previous findings that spatial variation in motility rates can change local aggregation and impact carrying capacity (Lu *et al*. 2020).

**Figure 5:**
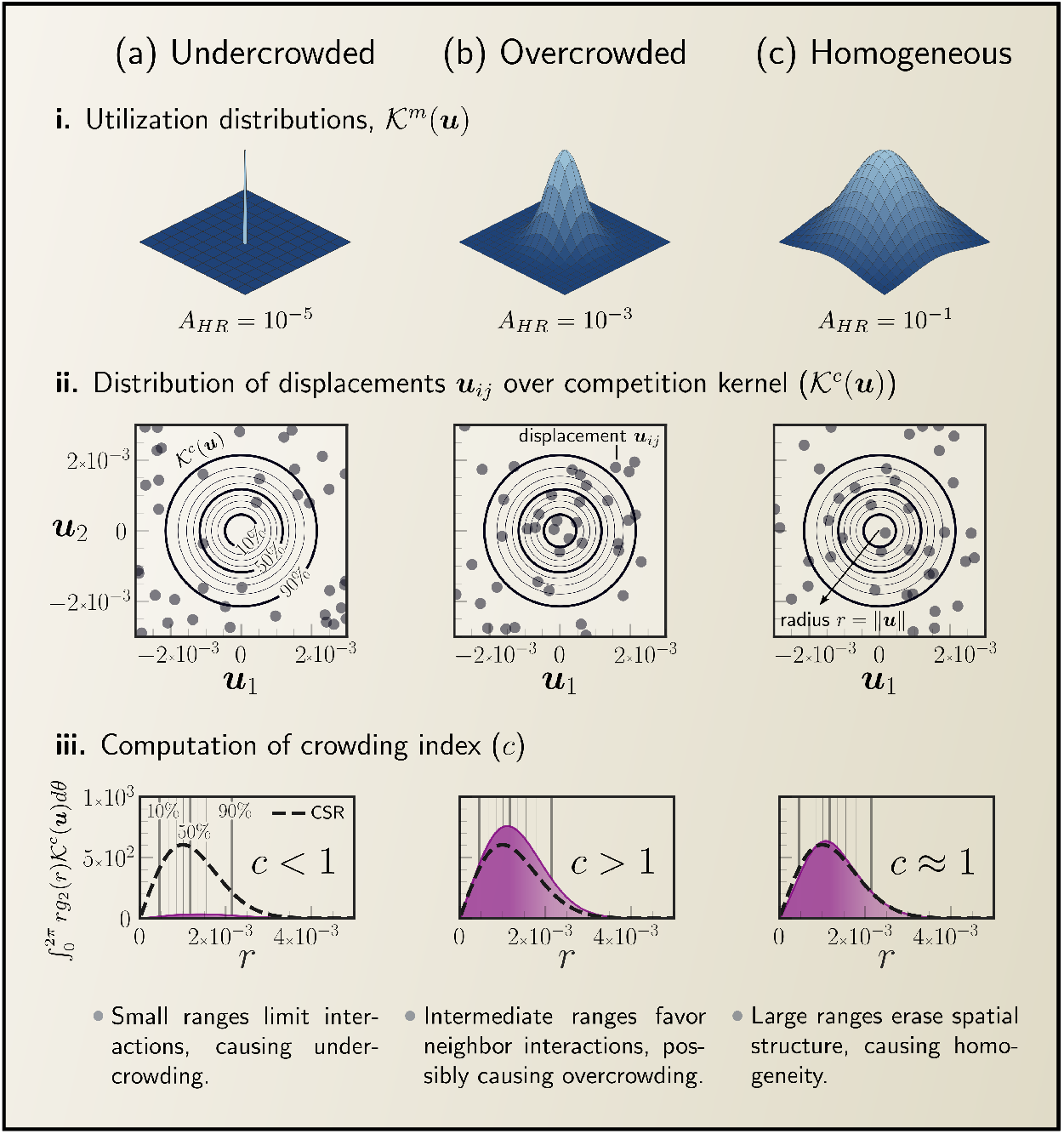
Movement behavior affects carrying capacity by influencing crowding—how much organisms are aggregated at the scale of competition. How organisms use space (utilization distributions; panel **i**) strongly influences the distribution of displacements among neighbors (***u***_*ij*_) relative to the competition kernel 𝒦^*c*^(***u***) (panel **ii**). The crowding index integrates the competition experienced at all competition scales (panel **iii**) to determine carrying capacity. (a) Small home ranges allow for spatial segregation, which limits interactions, resulting in undercrowding (*c <* 1) and increased abundances. (b) Intermediate home ranges lead to frequent interactions among neighbors, causing overcrowding (*c >* 1) and a higher risk of population collapse. (c) As organisms explore large portions of the available habitat, the population becomes homogeneous (*c* ≈ 1).

Organisms with very large ranges are homogeneously distributed in space, and interactions are well described by the law of mass action. Conversely, when home ranges are very small, organisms are effectively sessile and the population size is entirely determined by the competition and dispersal scales. Hence, this new framework captures two of the existing formulations of the logistic model as limits. It recovers the spatial logistic equations for sessile organisms when home ranges are vanishingly small (Bolker & Pacala 1997; Law *et al*. 2003) and Verhulst’s original formulation when home ranges are infinitely large (Verhulst 1838).

The simplicity of this new formalism allowed us to derive mathematical relationships that establish how home-range size and overlap determine intraspecific interactions and, ultimately, population size. Specifically, in the most realistic regime in which home-range crossing times are much shorter than the typical times between demographic events—so animal movement is statistically stationary within the organism’s lifetime (Fleming *et al*. 2014)—and the width of the competition kernel is negligible compared to the home-range size (Martinez-Garcia *et al*. 2020), intraspecific interactions are entirely determined by the long-term spatial pattern of individual space use. This is one of our key results, and it connects the strength of intraspecific interactions to movement parameters for which robust statistical estimators exist (Silva *et al*. 2022).

Leveraging this information about how 344 individual patterns of space use determine the strength of intraspecific interactions, we generalized the crowding index, originally developed for plant communities (Bolker & Pacala 1997; Wiegand *et al*. 2021), to motile organisms. This index identifies when range-resident animals can be treated as sessile organisms and predicts how spatial structure affects carrying capacity, showing that common home-range overlap metrics, such as the Bhattacharyya coefficient, quantify interaction strength inaccurately (Martinez-Garcia *et al*. 2020). Therefore, the crowding index links the mathematical framework of plant neighborhood models (Pacala & Silander 1985; Silander & Pacala 1985) to population and community dynamics in range-resident species. The original crowding index successfully integrated spatial information into demographic models of species-rich plant communities (Wiegand *et al*. 2021, 2025), and our generalization can be similarly applied to moving organisms, bridging movement and population ecology both theoretically and via novel statistical frameworks, the development of which is beyond the scope of this paper.

In models of population and community dynamics, crowding indexes derived from empirical or simulated spatial data rescale law-of-mass-action interaction rates, offering a straightforward way to account for movement behavior and spatial structure implicitly. This upscaling relies on two key assumptions: individual movement is statistically stationary over lifetimes—a common feature of many range-resident species (de Figueiredo *et al*. 2025; Fleming *et al*. 2014)—and individuals move independently of one another.

Connections with ecological data arise in two complementary directions. First, crowding indices can be estimated directly when both high-resolution movement data and competition scales are known. With only movement data, utilization distributions can approximate crowding by adapting estimators from spatial point pattern analysis (Baddeley *et al*. 2016; Wiegand *et al*. 2021), under the assumption of independent movement. These estimates allow researchers to quantify crowding in real populations and track such changes after interactions (Fagan *et al*. 2024). Comparing indices across populations of the same species could further provide a mechanistic measure of competition intensity, informing on how intraspecific interactions vary across environmental gradients (Olabarria *et al*. 2024).

Second, the range-resident logistic model can be embedded in hierarchical Bayesian frameworks that integrate movement and abundance data to jointly estimate demographic parameters, spatial-structure effects, and environmental covariates while propagating uncertainty across scales (Eisaguirre *et al*. 2025; Lu *et al*. 2020; McClintock *et al*. 2022). In these frameworks, observed counts are modeled as an observation process conditional on the true population size, which is treated as a latent variable. The range-resident logistic model can be used to model this latent population, conditioned on model parameters and the crowding index, thus helping the estimation of population density from spatial capture–recapture data when population size fluctuates due to movement and demography (McClintock *et al*. 2022).

The range-resident logistic model balances real-world complexity and a level of abstraction that allows for general insights. Beyond its most evident extension to a larger number of competing species and interactions, our framework can be refined by incorporating additional or alternative features of individual movement and life history traits.

First, incorporating variability in life history parameters (e.g. reproduction and death rates), alongside home range variability over time for a single individual and across individuals within the population (Ellison *et al*. 2020; Fleming *et al*. 2022) would allow investigations into how individual differences shape spatiotemporal population dynamics (Buchmann *et al*. 2011; Milles *et al*. 2020).

Second, the framework could be extended by including other realistic aspects of movement we omitted, such as perception, navigation, and memory (sensu Fagan *et al*. 2013; Nathan *et al*. 2008), transitions between behavioral modes (Blackwell 1997; Bläßle & Tyson 2016; Fleming *et al*. 2014; Smouse *et al*. 2010), or attraction/avoidance interactions (Surendran *et al*. 2019). This last aspect could show how complex behaviors, such as territory formation (Moorcroft & Lewis 2013; Potts & Lewis 2014) or range shifts following interaction (Fagan *et al*. 2024), impact carrying capacity.

Third, we explored the consequences of increased mortality rates in crowded populations, which can result from exhaustion of energetic reserves (Joshi & Mueller 1997) or an overall increase in stress levels (Gabriel *et al*. 2018; Pride 2005). Future model extensions could explore alternative population-regulation mechanisms, such as density-dependent reductions on birth rates (Lewis 2000; Plank & Law 2015) or environmental stochasticity (Doherty 1983). Accounting for birth-related regulation is particularly relevant if successful reproduction and offspring survival require exclusive territories and carrying capacity is determined by the extension of available territories (Ayllón *et al*. 2012; Carter *et al*. 2015; Hartmann *et al*. 2017; Knapton & Krebs 1974; López-Sepulcre & Kokko 2005).

Lastly, to focus on the role of home ranges, we used a phenomenological description of the dispersal process by employing a dispersal kernel that captures the distribution of natal dispersal displacements (Law *et al*. 2003; Nathan *et al*. 2012; Rogers *et al*. 2019). A promising avenue to expand the current framework is to incorporate more mechanistic descriptions of natal dispersal and habitat selection (de Oliveira *et al*. 2022; Mayor *et al*. 2009), while accounting for inter-organism variability in dispersal distance and its correlations with home range size (Bowman *et al*. 2002; Ronce *et al*. 1998).

In summary, we showed that accounting for range residency, a widespread feature of animal movement, is key to better understanding spatial patterns of aggregation, intensity of intraspecific interactions, and, ultimately, population dynamics. We established the theoretical foundation to formalize these relationships and provided mathematical expressions relating movement parameters, accessible from animal tracking data, to local crowding metrics and long-term population sizes in a single-species population dynamics model. This new framework, moreover, recovers previous formulations of the logistic model simply by manipulating home-range sizes, thereby providing a unifying theory for studying population dynamics in sessile, range-resident, and free-ranging species. Finally, because our theory balances ecological realism with mathematical tractability, it constitutes an important step toward the long-standing goal of bridging movement and population ecology.

## Acknowledgements

This study was financed, in part, by the São Paulo Research Foundation (FAPESP), Brasil - Process Number #2024/18255-0 (R.M.); the National Council for Scientific and Technological Development, Brazil - CNPq: 140096/2021-3 (R.M.); the Coordenação de Aperfeiçoamento de Pessoal de Nível Superior - Brasil (CAPES) - Finance Code 001 (R.M.); This work was partially funded by the Center for Advanced Systems Understanding (CASUS), which is financed by Germany’s Federal Ministry of Education and Research (BMBF) and by the Saxon Ministry for Science, Culture and Tourism (SMWK) with tax funds on the basis of the budget approved by the Saxon State Parliament; the São Paulo Research Foundation (FAPESP, Brazil) through BIOTA Young Investigator Research Grant No. 2019/05523-8 (R.M-G); ICTP-SAIFR grant no. 2021/14335-0 (R.M. and R.M.-G); the Simons Foundation through grant no. 284558FY19 (R.M-G). The National Science Foundation (NSF, USA) grant DBI_al_1915347 supported the involvement of J.M.C. and W.F.F. This research was supported by resources supplied by the Center for Scientific Computing (NCC/GridUNESP) of the São Paulo State University (UNESP). We thank Emilio Berti, Thorsten Wiegand and an anonymous reviewer for their constructive comments that helped improve the manuscript.

## Statement of authorship

RM and RM-G conceived the project and developed the model. RM performed all numerical and mathematical analyses with input from RM-G and PIP. RM and RM-G wrote the manuscript with input from all other authors. All authors discussed ideas and the results, and gave final approval for paper publication.

## Data accessibility statement

The code for individual-based simulations and analyses, along with the data and scripts used to generate the figures supporting this letter, are archived in the Zenodo repository: https://doi.org/10.5281/zenodo.15312822.

## Supporting Information

## A The crowding index for home range centers

In this appendix, we derive the expression for the crowding index based on home-range centers, Eq. (8), starting from its definition in terms of the instantaneous organism location, Eq. (5).

The position of a focal organism, *i*, and the displacement between this organism *i* and a neighbor *j*, both at time *t*, can be represented by two random variables 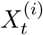 and 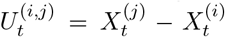 respectively. Each of these random variables has a probability density function (PDF), 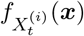 and 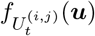, respectively. Similarly, we denote each individual’s home-range center as 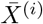, the displacement of the organism from its home range center at time *t* as 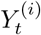, and the displacement between a pair of organisms’ home-range centers as 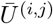. Using these definitions, the displacement between organisms is

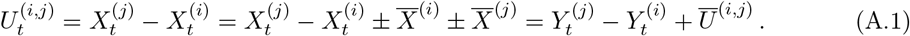

Because the PDF of a sum of random variables is the convolution of the PDFs of the random variables, the PDF of the displacement between organisms at time *t* is given by the convolution of PDF associated to 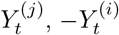 and 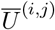. Mathematically, and using *f* to denote all PDFs, we can write this relationship as

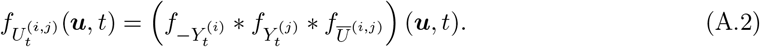

Additionally, the distribution of displacements between organisms *i* and *j*, averaged over all possible pairs of organisms, is directly related to the pair correlation function *g*_2_(***u***, *t*)

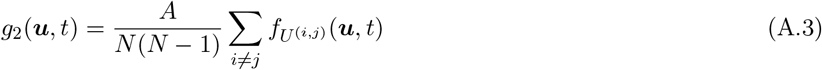

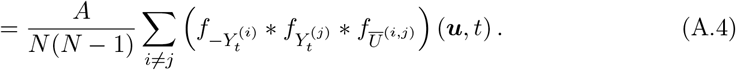

For range-resident organisms, the distribution of their position around its center tends to a constant asymptotic distribution directly related to its movement behavior, i.e. 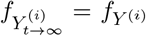 for any organism *i*. If we further assume that the home ranges of organisms are identical 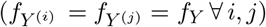 and symmetric (*f*_−*Y*_ = *f*_*Y*_), we can use the linearity of the convolution to write

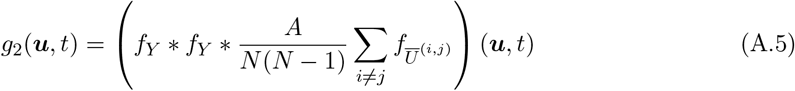

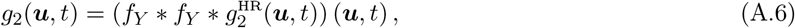

where we can identify 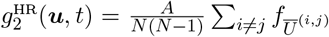 with the home-range center pair correlation function.

Finally, using Eq. (A.6), we can rewrite the crowding index as

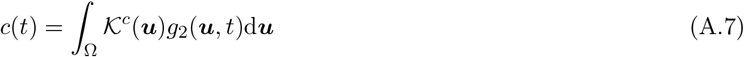

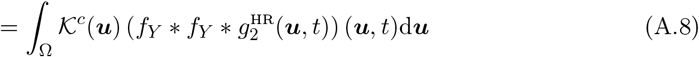

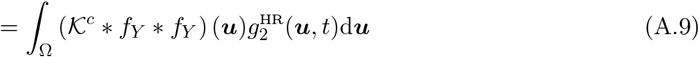

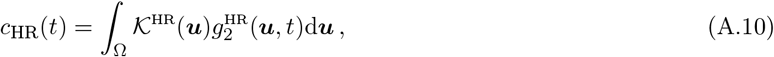

where we define the home-range competition kernel 𝒦^HR^(***u***) = (𝒦^*c*^ * *f*_*Y*_ * *f*_*Y*_) (***u***) and we denote the crowding index with the subscript HR to highlight that it is calculated using the distribution of HR centers instead of the using the spatial distribution of organisms.

## B Measurement of the crowding index using spatial information

This appendix explains how to measure the crowding index from simulation data. A simulation snapshot showing the spatial distribution of organisms can be described as a *N*-point pattern *P* = {***x***_*i*_} in which ***x***_*i*_ denotes the coordinates of the *i*-th point. Ignoring edge corrections because we are working with toroidal spatial domains, the pair correlation function of such point pattern is (Baddeley *et al*. 2016; Wiegand & Moloney 2014):

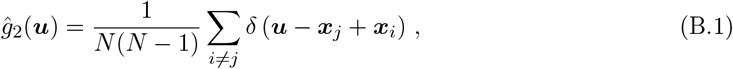

where *δ*(***u***) is the Dirac delta distribution and the summation is over all possible pairs of organisms. Substituting equation (B.1) into Eq. (8) we get

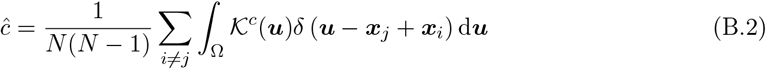

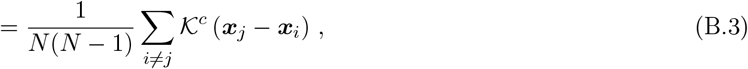

which can be used to measure the crowding index in the population, given a spatial distribution of organisms and for a known competition kernel.

Similarly, we apply the same procedure to compute the crowding index using the spatial pattern of home-range centers and space utilization functions. We first compute the home-range competition kernel 𝒦^HR^(***u***). Since the asymptotic displacement from the home-range center of the organisms following OU movement and the competition kernel 𝒦^*c*^ we implemented are Gaussian distributions centered at zero, their convolution is also a Gaussian distribution centered at zero. The crowding index based on home-range centers can then be computed as

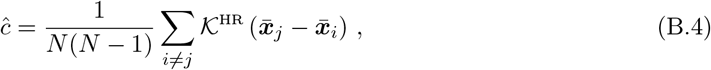

where 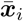 denotes the position of the home range center of the *i*-th organism.

We note that the derivation above allows for asymmetric, organism-specific, and pair-specific interaction kernels by taking 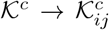. Thus, the proposed crowding index is compatible with asymmetric interactions, as seen in (Fagan *et al*. 2024), where the outcome of an encounter between two coyotes was asymmetric, leading to a one-sided shift in their home range overlap.

## C Supplementary Figures

**Figure C.1:**
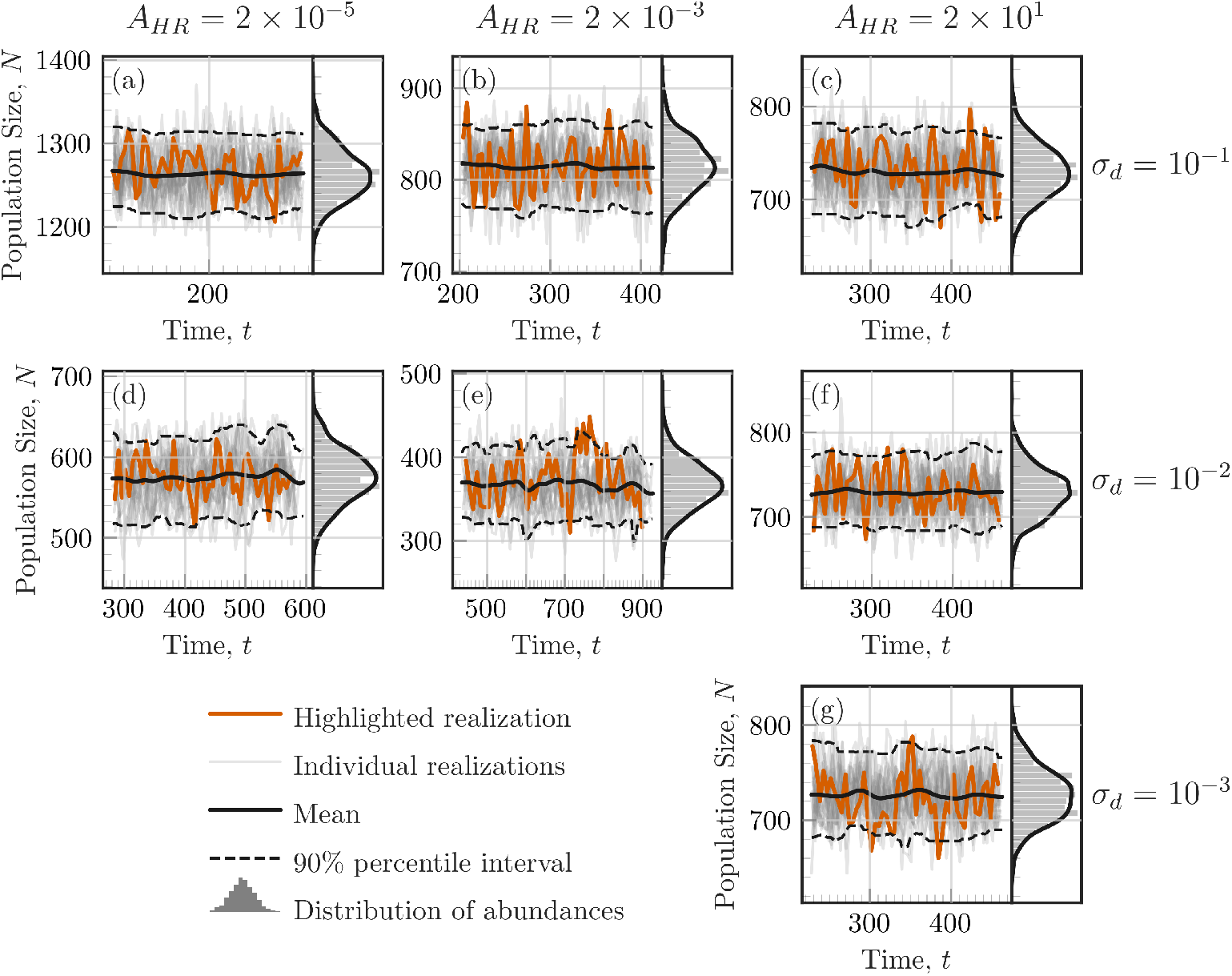
Diagnostic plots for stationarity of abundance. The abundance (y-axis) recorded in each realization as a function of simulation time (x-axis) is shown as light-gray lines. An arbitrary realization is highlighted in orange for easier visualization. The mean and 90% percentile interval of the abundance over 20 realizations are shown as full and dashed black lines, respectively. The distribution of recorded abundances over the measuring time (time range shown in x-axis of each panel) is shown as a histogram on the right. The data for this figure are excerpted from Fig. 2 on the main text.

**Figure C.2:**
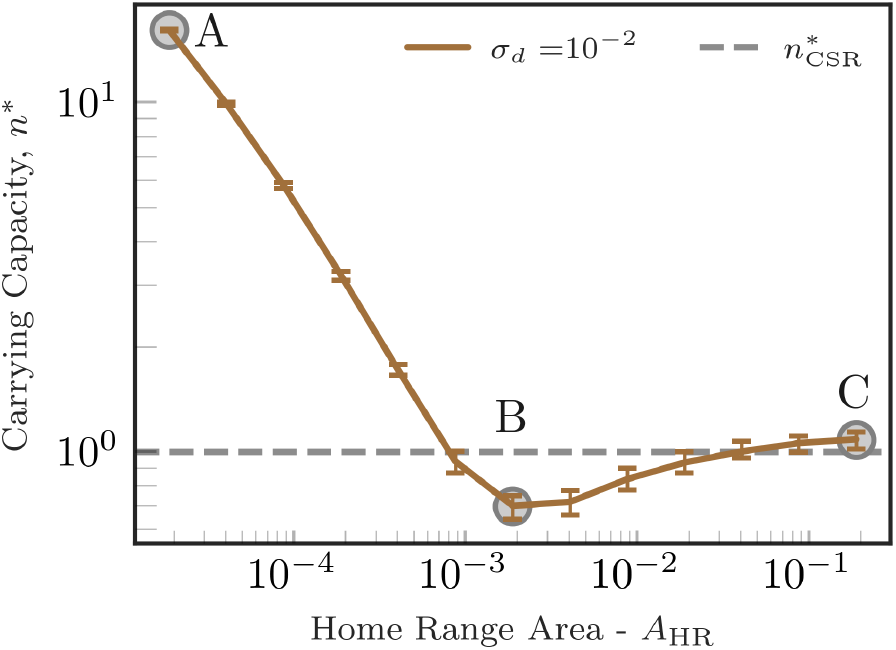
Home-range size alone can drive steady-state population sizes above or below the homogeneous carrying capacity. Increasing home-range size while all other spatial scales constant (dispersal and competition) can drive the stationary-state population size from much higher than predicted for a completely spatial random population, 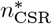 (A), to smaller (B), to finally match 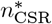 (C). Standard deviations of competition and dispersal kernels were set to *σ*_*q*_ = 10^−3^ and *σ*_*d*_ = 10^−2^, respectivelly.

**Figure C.3:**
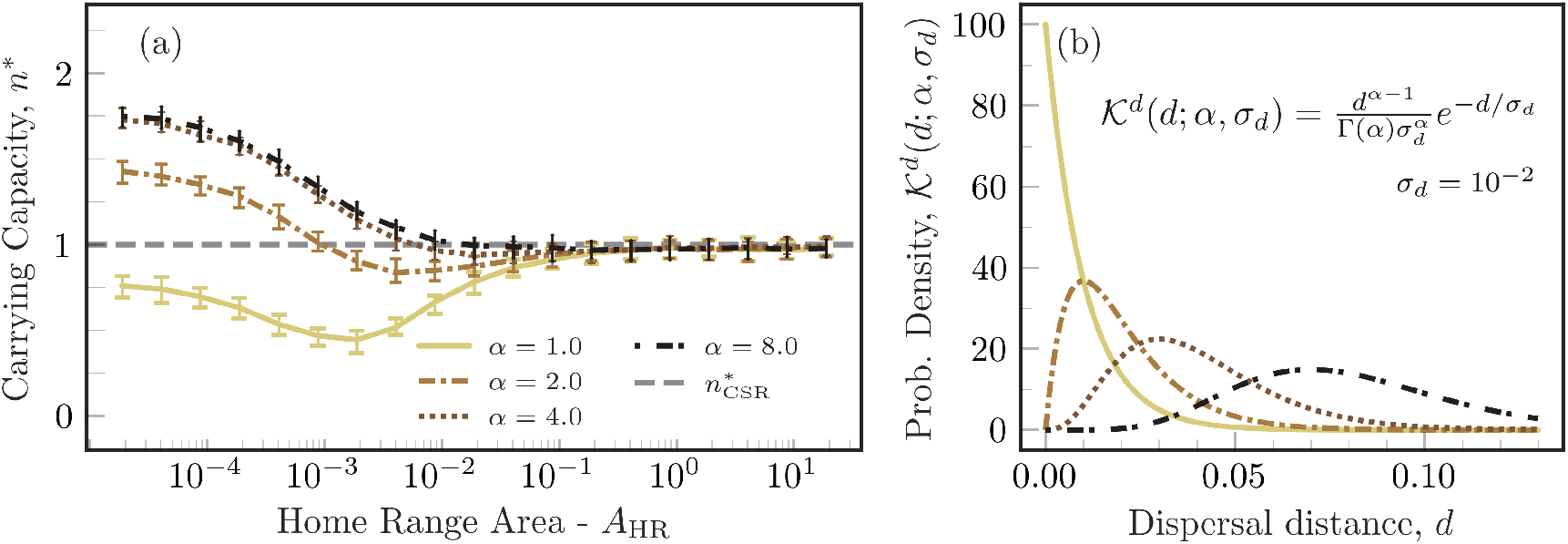
Carrying capacity versus home-range size for gamma-distributed dispersal distances. The results obtained by varying the shape parameter of the gamma distribution mirror those obtained by varying the standard deviation of the bivariate normal distribution. For this analysis, the dispersal kernel was defined such that the radial distance from the parent’s home range center follows a gamma distribution with shape parameter *α* ∈ [1, 2, 4, 8] (different colors and line styles as indicated in the legend of panel (a)) and constant dispersal and competition scale parameters *σ*_*d*_ = *σ*_*q*_ = 10^−2^. The direction of dispersal was assumed to be uniformly distributed in [0, 2*π*). (a) Carrying capacity measured in the simulations and scaled by the carrying capacity of the non-spatial model, as in Fig. 2 of the main text. (b) Dispersal probability density function for the dispersal kernels used in (a). Color lines correspond to values of *α* as indicated in the legend of panel (a). The dispersal and competition scales remained constant for all simulations, *σ*_*d*_ = *σ*_*q*_ = 10^−2^.

**Figure C.4:**
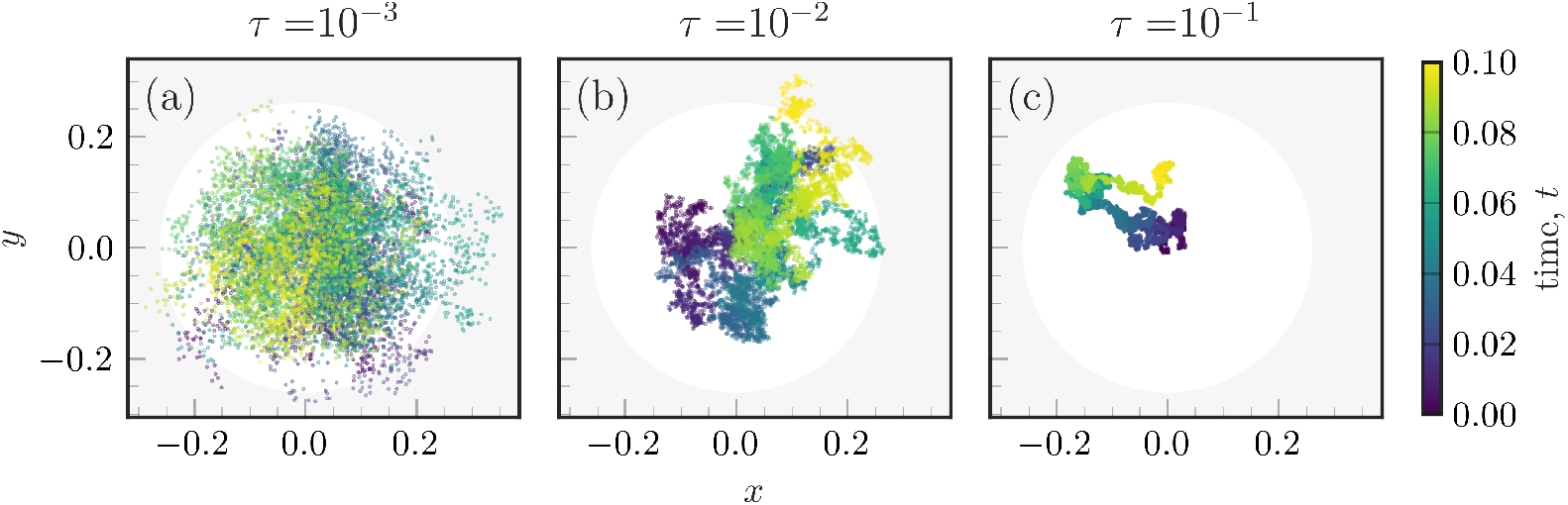
The effect of home range crossing time on how fast OU-moving individuals cover their entire home range. Organisms take longer to explore their entire home range when the home-range crossing time *τ* increases (from panel a to c), keeping home-range size fixed, *A*_HR_ = 0.20(*a*.*u*.). In the three panels, we simulated a single trajectory for a total of *t* = 0.1(*a*.*u*.) with time increments of *dt* = 10^−5^(*a*.*u*.). For smaller values of *τ* (panel a), the organism effectively explores the entire HR. However, if *τ* increases but we keep the simulation time constant, the organism explores a more restricted region of its home range. In the limit where *τ* tends to zero, the trajectory loses all the autocorrelation between consecutive locations, which becomes an independent and identically distributed variable (IID). Consequently, the organism can be found anywhere within the home range at each time step.

**Figure C.5:**
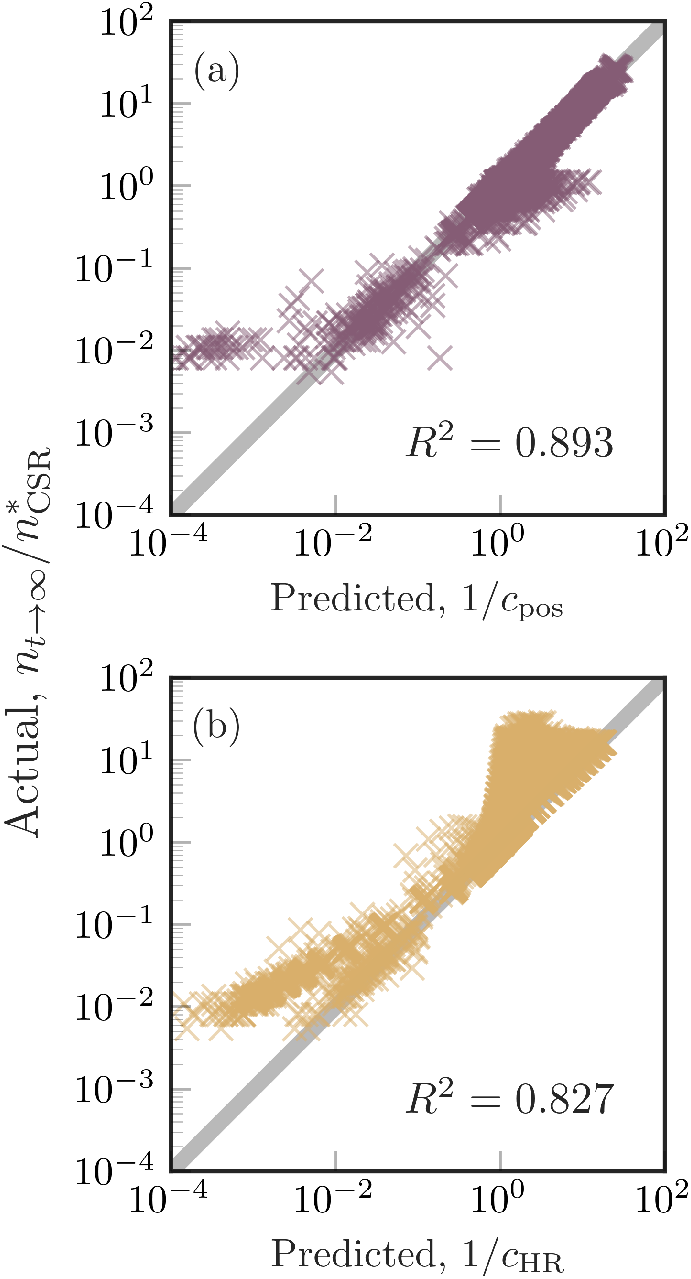
Accuracy of the crowding index for broader dataset. The crowding index measured using only information about spatial scales across all the simulations matches the measurement of the carrying capacity of the population. The abundance of the population in the simulation normalized by the carrying capacity of a homogeneous population (y-axis) is compared to the inverse of the crowding coefficient (x-axis). (a) The crowding index was measured using the actual position of the organisms as the average local density experienced by each organism in a snapshot of the simulation. (b) The crowding index was measured based on the expected mortality rate an organism experiences given its home-range size and the distances between its home range center and that of its neighbors. The complete agreement (1-1 line) is represented as a gray line in the background of each panel and the *R*^2^ is reported as an indicative of the goodness of fit. No data was filtered out of the analysis, apart from simulations in which extinction was observed.

